# Temporally distinct oscillatory codes of retention and manipulation of verbal working memory

**DOI:** 10.1101/2021.03.13.435253

**Authors:** Yuri G. Pavlov, Boris Kotchoubey

## Abstract

Most psychophysiological studies of working memory (WM) target only the short-term memory construct, while short-term memory is only a part of the WM responsible for the storage of sensory information. Here, we aimed to further investigate oscillatory brain mechanisms supporting the executive components of WM – the part responsible for the manipulation of information. We conducted an exploratory reanalysis of a previously published EEG dataset where 156 participants (82 females) performed tasks requiring either simple retention or retention and manipulation of verbal information in WM. A relatively long delay period (>6s) was employed to investigate the temporal trajectory of the oscillatory brain activity. Compared to baseline, theta activity was significantly enhanced during encoding and the delay period. Alpha-band power decreased during encoding and switched to an increase in the first part of the delay before returning to the baseline in the second part; beta-band power remained below baseline during encoding and the delay. The difference between the manipulation and retention tasks in spectral power had diverse temporal trajectories in different frequency bands. The difference maintained over encoding and the first part of the delay in theta, during the first part of the delay in beta, and during the whole delay period in alpha. Our results suggest that task-related modulations in theta power co-vary with the demands on the executive control network; beta suppression during mental manipulation can be related to the activation of motor networks; alpha is likely to reflect the activation of language areas simultaneously with sensory input blockade.

## Introduction

Many typical WM tasks, in which the relationships with cortical oscillations are investigated, contain an initial period of encoding, followed by a period of maintenance, i.e., the delay period. Whereas in behavioral experiments delay periods vary in a wide range, from less than a second to tens of seconds (Berman, Jonides, & Lewis, 2009; Lewandowsky, Oberauer, & Brown, 2009; Oberauer et al., 2018), in neurophysiological studies the duration of delay is limited and usually depends on the temporal resolution of the neurophysiological measurements. Thus, the typical delay duration in EEG studies is less than 3 seconds (Pavlov & Kotchoubey, 2021), while fMRI studies employ relatively long delay periods of 6 seconds or longer. Using only short delay periods typical for EEG research may complicate the comparison with the results obtained in numerous behavioral studies. The present research takes advantage of the high temporal resolution in EEG recordings to expand our knowledge of WM delay activity on different time scales.

Behavioral research in healthy volunteers largely indicates a small (or even zero) effect of the delay duration on verbal WM performance when rehearsal is not suppressed (Oberauer, Farrell, Jarrold, & Lewandowsky, 2016; Oberauer et al., 2018). On the one hand, oscillatory brain activity during the delay period is unlikely to be static although the evidence for the opposite is not overwhelming. EEG studies with typical delay periods (<3s) clearly demonstrated sustained alpha during delay (Bashivan, Bidelman, & Yeasin, 2014; Hsieh, Ekstrom, & Ranganath, 2011; Hu et al., 2019; Jensen, Gelfand, Kounios, & Lisman, 2002; Roberts, Hsieh, & Ranganath, 2013; Tuladhar et al., 2007). However, a visual WM study with EEG recordings used a 6 s long delay period and found a dissociation between its early and late parts (Ellmore, Ng, & Reichert, 2017). The early delay was characterized by a prevalence of alpha activity, but after about 4 s alpha completely disappeared, marking the onset of the late delay phase. Verbal WM studies did not test explicitly the spectral power changes during the long delay.

If brain oscillations are indeed critical for the maintenance of information in WM one might expect a sustained pattern of activity in all relevant frequency bands for the whole duration of the delay period. For example, assuming that alpha reflects the engagement or active inhibition of attention (e.g., Poch et al., 2014; Sauseng et al., 2005; Schneider et al., 2019), the fluctuation of attention during the delay may be expected but the overall level of activity should differ from the baseline level. Furthermore, if theta activity plays a major role in executive functioning (Cavanagh & Frank, 2014; Kawasaki, Kitajo, & Yamaguchi, 2010) or storage of temporal order (Hsieh & Ranganath, 2014), then both functions are required up to the end of the delay period. If, on the other hand, oscillatory brain activity during the delay is of transient nature, this would affect the interpretation of the role of oscillations in memory. Thus, given the absence of the effect of duration of the delay on behavioral performance (e.g., accuracy or reaction time), one may suggest that different neurophysiological mechanisms can underlie similar behavioral outcomes.

A core feature of the verbal WM construct is the requirement to retain and manipulate information simultaneously (Oberauer, Süß, Schulze, Wilhelm, & Wittmann, 2000). The manipulation abilities allow to mentally reorganize information to perform everyday tasks such as reading, mental math, navigation in the traffic, updating information during a conversation. Despite its importance, neural correlates of the WM manipulations and associated increased demands on the executive control networks have been a topic of interest in a small number of EEG studies. Available research points out the role of theta activity in the manipulation of information in WM (Berger, Omer, Minarik, Sterr, & Sauseng, 2014; Griesmayr, Gruber, Klimesch, & Sauseng, 2010; Itthipuripat, Wessel, & Aron, 2013; Kawasaki et al., 2010; Pavlov & Kotchoubey, 2017) but other rhythms have received much less attention. Here, we expected to provide a more complete picture on the spectral fingerprints of the executive control of WM by tracking the temporal dynamics of the difference between the simple retention and manipulation tasks.

In the current study, we used our previously published dataset (Pavlov & Kotchoubey, 2020) where we recorded EEG in a WM task that engaged the manipulation and retention functions of WM. While in the previous publication we tested hypotheses concerning the relationship between individual differences in WM and brain oscillations, here we performed exploratory analyses of general effects. We aimed (1) to replicate, in a large sample, previously established findings such as an increase of theta activity with increasing demand on executive control elicited by increasing WM load and the task to manipulate information in WM, while providing new results on how these conditions affect alpha and beta activity; (2) to generalize these findings to a less frequently employed complex item-location task; and (3) to test how the extant data, typically obtained in experiments with short delay periods, generalize to a task requiring retention or manipulation of information for a rather long time interval (>6 s), comparable with typical delay periods in behavioral and fMRI experiments. In particular, we tested whether the typical continuous increase in theta and alpha activity during delay shows the same pattern over a longer period of time.

## Methods

All data obtained in the study, including raw and preprocessed EEG recordings, are organized in BIDS format and freely available (doi: 10.18112/openneuro.ds003655.v1.0.0). Below, we describe essential methodological aspects but we refer to the original article where the dataset was published first for details (Pavlov & Kotchoubey, 2020).

### Participants

156 participants (82 females, mean age = 21.23, SD=3.22) comprised the final sample. The participants had normal or corrected-to-normal vision and did not report any history of neurological or mental diseases. All of them were Russian native speakers. The experimental protocol was approved by the Ural Federal University ethics committee.

### Task

Our task and behavioral results are described in detail elsewhere (Pavlov & Kotchoubey, 2020). Briefly, we used a task with temporally distinct encoding and maintenance processing stages (see Figure 1). The experiment entailed six different conditions: maintenance in memory of 5, 6 or 7 simultaneously presented letters in the alphabetical (manipulation task) or forward (retention task) order. In the retention task the participants had to maintain in memory the original set as it was presented, and in the manipulation task they had to, first, mentally reorganize the letters into the alphabetical order and then maintain the result in memory. After 6.7 s delay, a letter-digit probe appeared and the participants indicated whether the probe was on the corresponding position either in the original set (retention task), or in the set resulted from the alphabetical reordering (manipulation task). Each of the six conditions (retention or manipulation of 5, 6 or 7 letters sets) entailed 20 consecutive trials. These six blocks of 20 trials were presented in a random order.

**Figure 1.**
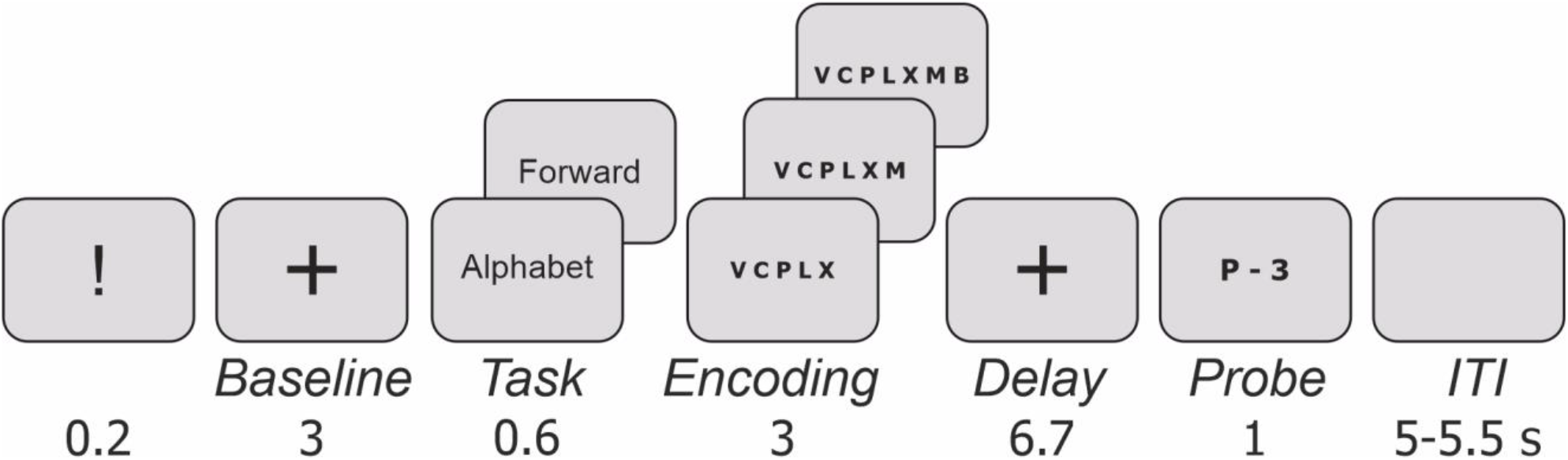
The experimental paradigm. Sets of Russian alphabet letters (5, 6, or 7) written in capitals were used as stimuli. The letters had been selected randomly from the alphabet, had random order, and no repetitions in the sets. An analogue using Latin letters and English words is shown. ITI – inter-trial interval.

### Electroencephalography

The EEG was recorded from 19 electrodes arranged according to the 10-20 system using Mitsar-EEG-202 amplifier with averaged earlobes reference. Two additional electrodes were used for horizontal and vertical EOG. EEG data were acquired with 500 Hz sampling rate and 150 Hz low-pass filter. For further processing, 1 Hz high-pass, 45 Hz low-pass and 50 Hz notch offline filters were applied with the EEGLAB firfilt function with default settings.

The procedure of EEG artifacts suppression and removal was conducted in two steps. At the first step, in order to suppress ocular activity artifacts, Independent Component Analysis (ICA) was performed using AMICA algorithm with default settings (Palmer, Kreutz-Delgado, & Makeig, 2012). The components clearly related to blinks and eye movements were identified and removed after visual exploration of the data. 3.32 components rejected on average (median: 3, range: 1-7). Then, epochs in the [-14200 2200 ms] interval where 0 is the onset of the probe were created. The epochs still containing artefacts were visually identified and discarded. EEGLAB toolbox (Delorme & Makeig, 2004) for MATLAB was used for the data preprocessing.

Time-frequency analysis was performed on the preprocessed single trial data between 1 and 45 Hz with 1 Hz steps using Morlet wavelets with the number of cycles varying from 3 to 12 in 45 logarithmically spaced steps for each participant and condition, separately. The analysis time window was shifted in steps of 10 ms. Spectral power was baseline-normalized by computing the percent change of the power in respect to the [-11500 to -10500] ms time interval, which corresponded to one second of the baseline fixation from -1200 to -200 ms before presentation of the task (see Figure 1). The time-frequency analysis was performed by means of the Fieldtrip toolbox (Oostenveld, Fries, Maris, & Schoffelen, 2011).

### Statistics

In order to decrease the number of factors employed in statistical calculations, we defined frequency-channels regions of interest by, first, selecting three frequency bands showing a response during both encoding and delay periods, and, second, by identifing physiologically meaningful and supported by the literature symmetric areas (groups of channels) with strongest manifestaton of the respective oscillations. Thus, theta (4-8 Hz) had the maximal power at Fz, alpha (9-14 Hz) at posterior channels (T5, P3, O1, T6, P4, O2) and beta (16-22 Hz) at central channels (C3, Cz, C4). The ROIs were defined on the basis of visual inspection of the grand average over all conditions.

For the statistical analyzes described in the following sections, the EEG power in theta, alpha and beta frequency bands was averaged in three time intervals. The first time interval corresponded to the presentation of the stimuli (2500 ms starting from 500 ms after the onset of the memory sets, see Figures 2, 3, and 4). This interval will be referred to as Encoding. Then, the segment of 6000 ms corresponding to the delay period starting from -6000 ms to 0 ms (where 0 is the onset of the probe) was divided into two time intervals: Delay1 (from -6000 ms to -3000 ms) and Delay2 (from -3000 ms to 0 ms). We excluded the initial 700 ms from the onset of Delay period to reduce the effect of the evoked response activity distorting frequency data (Babu Henry Samuel, Wang, Hu, & Ding, 2018; Ikkai, Blacker, Lakshmanan, Ewen, & Courtney, 2014; van Gerven, Bahramisharif, Heskes, & Jensen, 2009).

**Figure 2.**
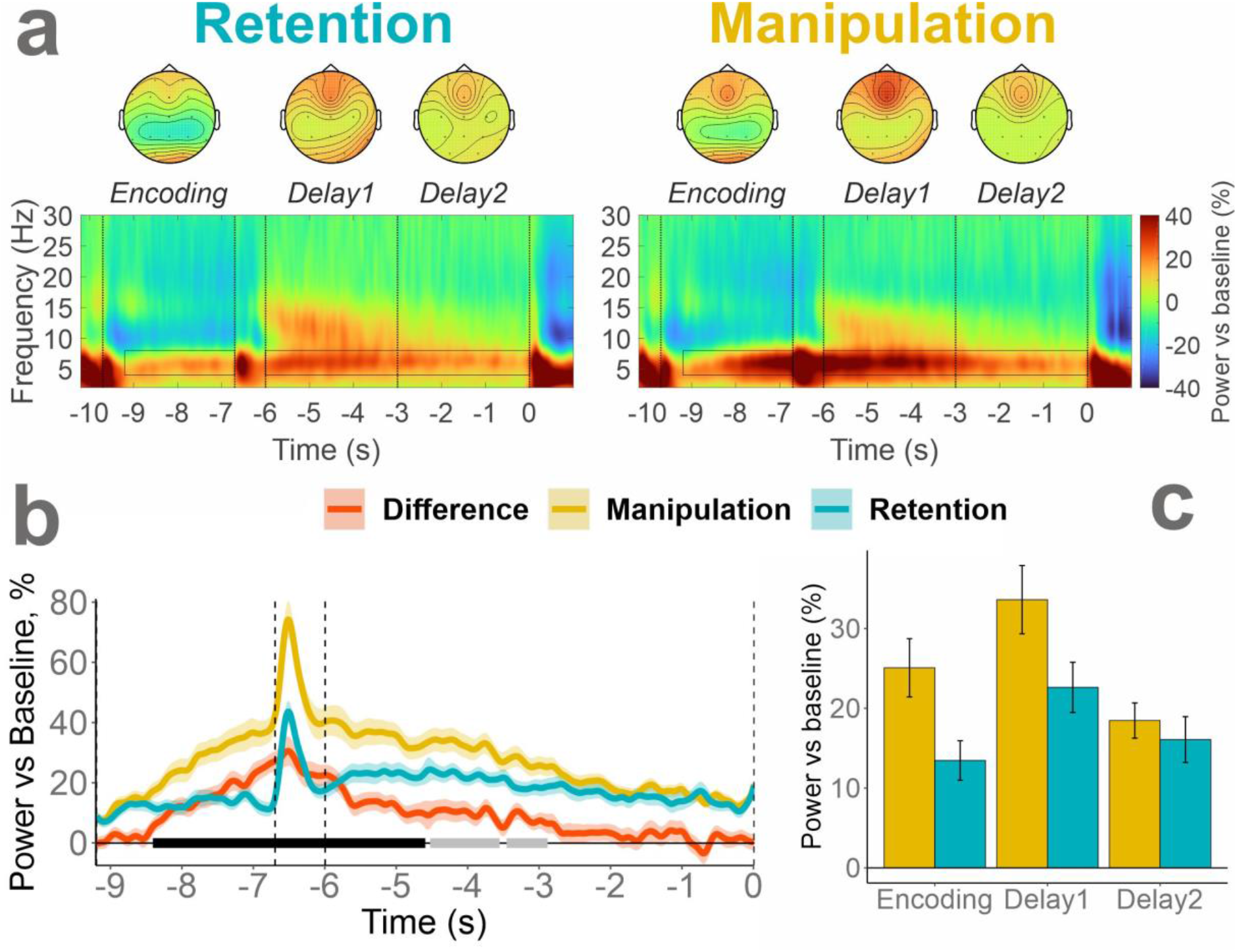
Task and time interval effects in the theta frequency band at Fz. (a) Time-frequency maps in manipulation and retention tasks. Horizontal boxes mark time-frequency windows of interest (4-8 Hz, last 2.5 s of the encoding, last 6 s of the delay). Vertical bars mark the encoding, the first and second parts of the delay period. The topographical maps show the distribution of the spectral power in the corresponding time intervals and tasks. (b) The time dynamics of theta activity in Retention, Manipulation tasks and their difference. Cluster-based permutation test results are marked with black (p<0.005) and grey (p<0.05) bars along the baseline. (c) Bar plot of the Time interval by Task interaction. The shading (b) and error bars (c) show the standard errors of the mean.

**Figure 3.**
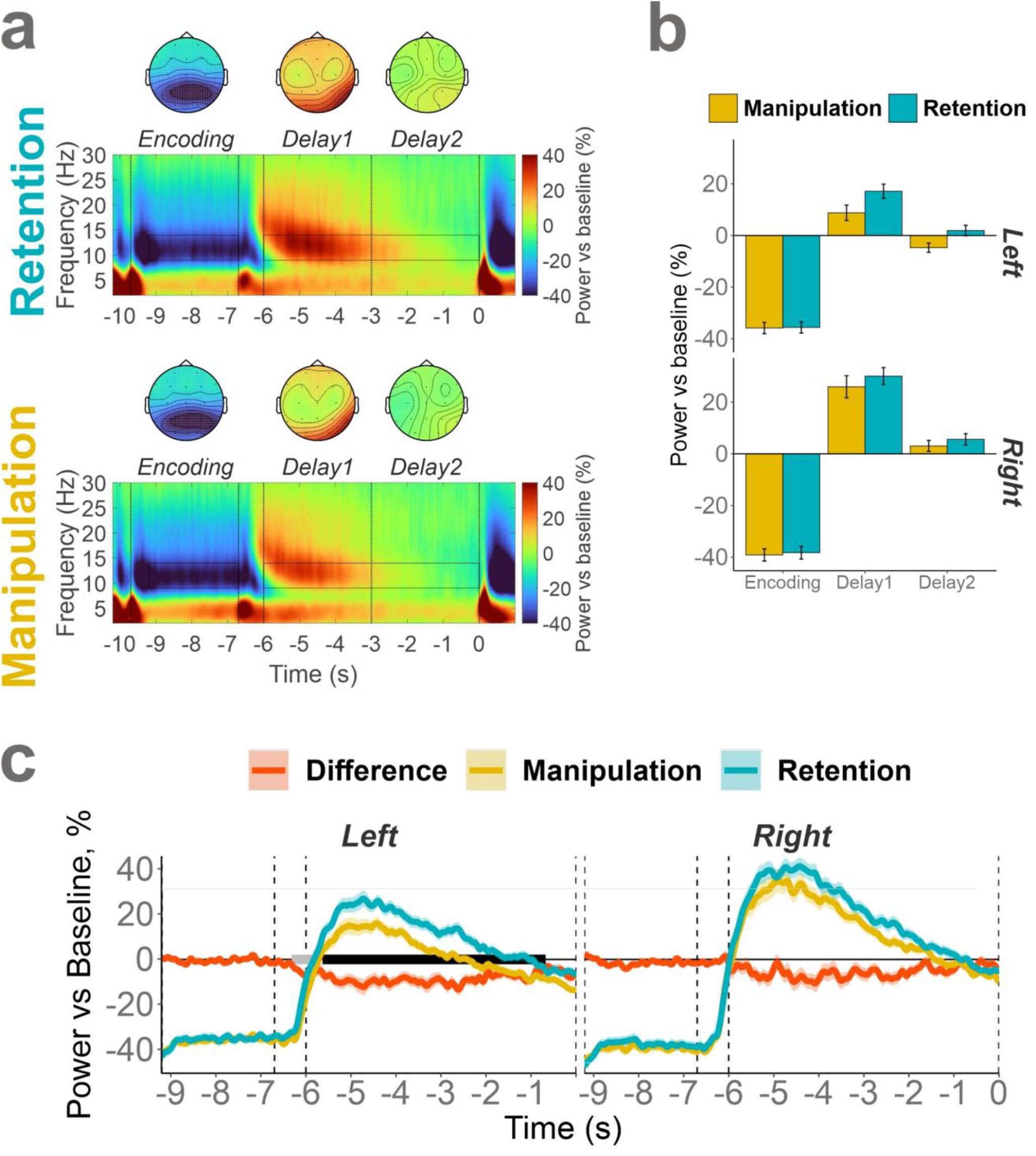
The task, time interval, and Hemisphere effects in the alpha frequency band over posterior ROI (T5, P3, O1, T6, P4, O2). (a) Time-frequency maps in manipulation and retention tasks. Horizontal boxes mark time-frequency windows of interest (9-14 Hz, last 2.5 s of the encoding, last 6 s of the delay). The vertical bars mark the encoding, the first and second parts of the delay period. The topographical maps show the distribution of the spectral power in the corresponding time intervals and tasks. (b) Bar plot of the Time interval by Task by Hemisphere interaction. (c) The time dynamics of alpha activity in Retention, Manipulation tasks and their difference in the left (T5, P3, O1) and right (T6, P4, O2) hemisphere channels. Cluster-based permutation test results are marked with black (p<0.005) and grey (p<0.05) bars along the baseline. No significant clusters were found in the right hemisphere. The shading (c) and error bars (b) show the standard errors of the mean.

**Figure 4.**
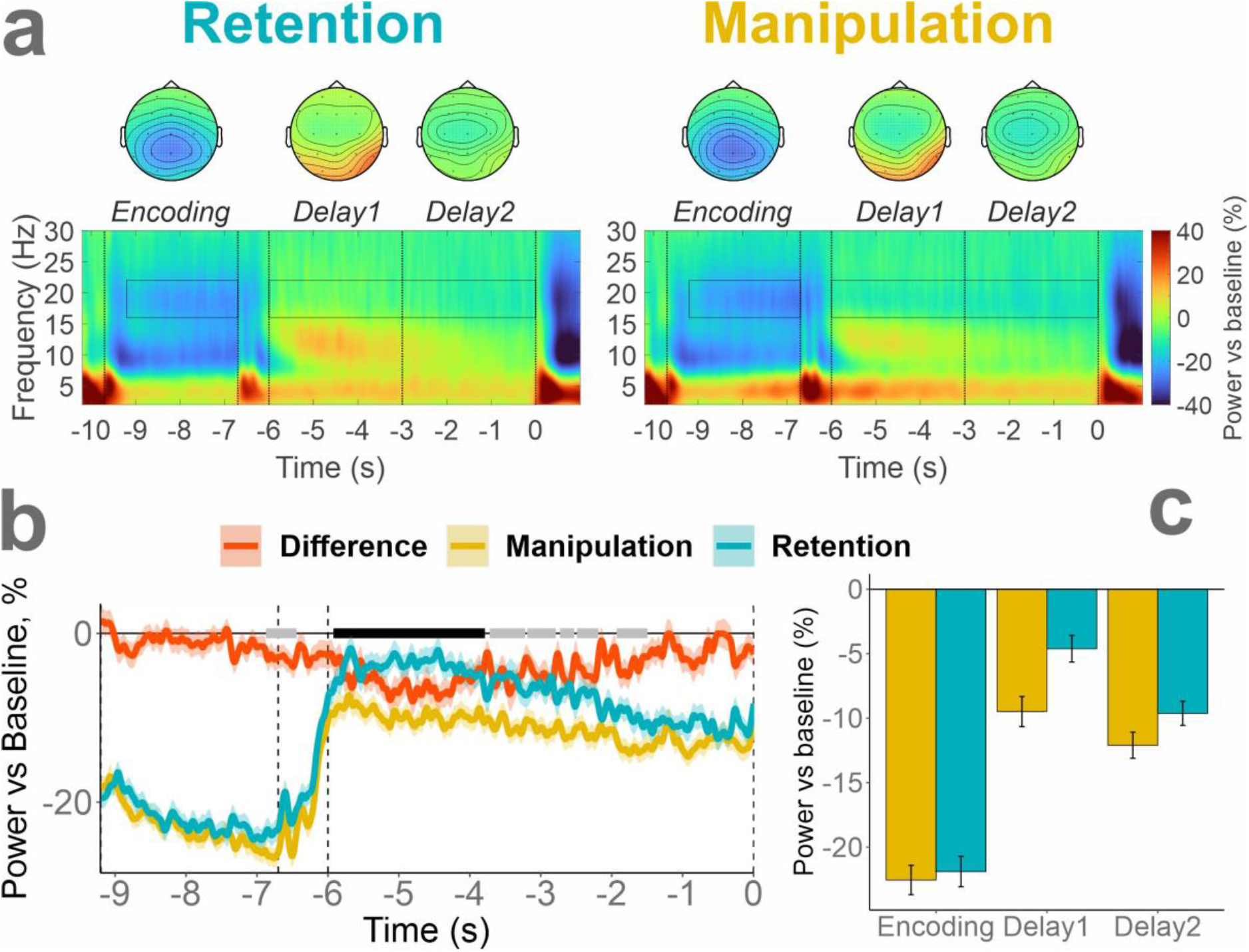
Task and time interval effects in the beta frequency band over central ROI (C3, Cz, C4 channels). (a) Time-frequency maps in manipulation and retention tasks. The black boxes mark time-frequency windows of interest (16-22 Hz, last 2.5 s of the encoding, last 6 s of the delay). The vertical bars mark the encoding, the first and second parts of the delay period. The topographical maps show the distribution of the spectral power in the corresponding time intervals and tasks. (b) The time dynamics of beta activity in Retention, Manipulation tasks and their difference. Cluster-based permutation test results are marked with black (p<0.005) and grey (p<0.05) bars along the baseline. (c) Bar plot of the Time interval by Task interaction. The shading (b) and error bars (c) show the standard errors of the mean.

In order to explore the effects of experimental conditions on EEG spectral power, a repeated-measures (RM) ANOVA was carried out with within-subject factors Task (2 levels: Retention or Manipulation), Load (5, 6 or 7 letters to memorize), and Time interval (TimeInt for short; with 3 levels: Encoding, Delay1 and Delay2). The analysis was conducted separately for theta and beta frequency bands. To analyze alpha activity an additional factor Hemisphere (2 levels: Left (P3, T5, O1), Right (P4, T6, O2)) was used.

Simple effects in ANOVA interactions were further explored with paired t-tests. The Holm method for correction for multiple comparisons was used where applicable. The alpha level was set to 0.005. In the following ANOVAs, asterisks will be used to highlight the effects of different magnitude with partial eta squared equal to or larger than .0099*, .0588**, and .1379*** as benchmarks for small*, medium**, and large*** effect sizes, respectively (Cohen, 1992; Richardson, 2011). The effect size is marked in the statistical output tables only in the case of statistical significance (p<0.005). In addition to the ANOVAs, and to confirm the conclusions derived from them, we performed cluster-based permutation tests (*clusterlm* function with 5000 permutations) using *permuco* package for *R* (Frossard & Renaud, 2019).

All statistical calculations were performed in *R*, version 3.6.3 (R Core Team, 2013).

## Results

### Behavior

As was reported in our previous publication in detail, accuracy was significantly lower in the manipulation task than in the retention task, and monotonically decreased with load. Importantly, the accuracy was nearly identical during manipulation of 5 letters and retention of 7 letters. The accuracy data is reported in Table 1 below.

**Table 1.** Descriptive statistics for accuracy

| *Load* | *Task* | *Mean % correct responses* | *SD* |
| --- | --- | --- | --- |
| 5 | Manipulation | 79.55 | 14.33 |
| 5 | Retention | 92.28 | 9.02 |
| 6 | Manipulation | 66.22 | 15.51 |
| 6 | Retention | 88.65 | 10.94 |
| 7 | Manipulation | 65.61 | 13.52 |
| 7 | Retention | 80.71 | 13.55 |

Notes: *SD* – standard deviation

### Theta

Theta activity changed as a function of time interval (main effect of TimeInt, see Figure 2, and Table 2 for statistical output). Specifically, across both tasks, the relative theta power during Encoding was lower than during Delay1 (t(155) = 7.34, p < 0.001, dz = 0.59) but was not significantly different from Delay2 (t(155) = 1.09, p = 0.27, dz = 0.09). The increase of theta in the first part of the Delay was stronger than in the second part (t(155) = 4.99, p < 0.001, dz = 0.40).

**Table 2.** ANOVA RM statistics in theta

| | *DF* | *F* | $\eta^2$ | *p* |
| --- | --- | --- | --- | --- |
| **TimeInt** | 2, 298 | 22.69 | 0.128 | **< 0.001 \*\*** |
| **Task** | 1, 155 | 17.16 | 0.100 | **< 0.001 \*\*** |
| **Load** | 2, 275 | 1.09 | 0.007 | 0.333 |
| **TimeInt x Task** | 2, 249 | 9.52 | 0.058 | **< 0.001 \*** |
| TimeInt x Load | 2, 341 | 1.45 | 0.009 | 0.236 |
| Task x Load | 2, 249 | 4.58 | 0.029 | 0.017 |
| TimeInt x Task x Load | 2, 326 | 1.85 | 0.012 | 0.157 |

\* small \*\* medium \*\*\* large effect size and $p < 0.005$

The theta increase was stronger in the manipulation as compared to the retention task (main effect of Task). This task effect was present during encoding (t(155) = 4.96, p < 0.001, dz = 0.40) and Delay1 (t(155) = 4.28, p < 0.001, dz = 0.34) but did not attain significance during Delay2 (t(155) = 0.99, p = 0.33, dz = 0.08), resulting in a significant TimeInt x Task interaction.

### Alpha

As illustrated in Figure 3, alpha activity was strongly affected by the time interval (see Table 3 for statistical output). After an initial suppression during Encoding the pattern reversed in Delay1. In Delay2, the relative alpha power was not significantly different from the baseline level (t(155) = 0.84, p = 0.4, dz = 0.07). As a result, alpha in Delay1 was significantly larger than in Delay2 (t(155) = 8.504, p < 0.001, dz = 0.68).

**Table 3.** Alpha ANOVA RM

| | *DF* | *F* | $\eta^2$ | *p* |
| --- | --- | --- | --- | --- |
| **Hemisphere** | 1, 155 | 39.55 | 0.203 | **< 0.001\*\*\*** |
| **TimeInt** | 2, 263 | 222.36 | 0.589 | **< 0.001\*\*\*** |
| Task | 1, 155 | 7.66 | 0.047 | 0.006 |
| Load | 2, 304 | 0.73 | 0.005 | 0.479 |
| **Hemisphere x TimeInt** | 1, 222 | 64.15 | 0.293 | **< 0.001\*\*\*** |
| Hemisphere x Task | 1, 155 | 4.69 | 0.029 | 0.032 |
| TimeInt x Task | 2, 246 | 5.55 | 0.035 | 0.008 |
| Hemisphere x Load | 2, 304 | 0.09 | 0.001 | 0.909 |
| TimeInt x Load | 3, 489 | 3.76 | 0.024 | 0.010 |
| Task x Load | 2, 301 | 2.22 | 0.014 | 0.112 |
| **Hemisphere x TimeInt x Task** | 2, 240 | 7.18 | 0.044 | **0.002 \*** |
| Hemisphere x TimeInt x Load | 3, 490 | 1.43 | 0.009 | 0.230 |
| Hemisphere x Task x Load | 2, 307 | 0.19 | 0.001 | 0.829 |
| TimeInt x Task x Load | 3, 500 | 0.55 | 0.004 | 0.660 |
| Hemisphere x TimeInt x Task x Load | 3, 421 | 0.82 | 0.005 | 0.472 |

\* *small* \*\* *medium* \*\*\* *large effect size and $p < 0.005$*

On average, the relative alpha power was larger over the right compared to the left hemisphere (main effect of Hemisphere). This effect was modulated by Time interval (TimeInt x Hemisphere interaction, see Figure 3b and c). The difference between the alpha power in the left and right hemispheres during encoding (t(155) = 3.18, p = 0.002, dz = 0.26) was smaller than during the first (t(155) = 8.36, p < 0.001, dz = 0.67) and second part of delay (t(155) = 5.57, p < 0.001, dz = 0.48).

The three ANOVAs with two levels of TimeInt factor resulted in significant TimeInt x Hemisphere interactions (Encoding and Delay1: F(1, 155) = 79.9, p < 0.001, η^2^ = 0.34; Encoding and Delay2: F(1, 155) = 45.3, p < 0.001, η^2^ = 0.23; Delay 1 and Delay 2: F(1, 155) = 49, p < 0.001, η^2^ = 0.24), confirming that all three time intervals differed in respect of the alpha activity asymmetry (see Figure 3).

Although both alpha increase (delay) and suppression (encoding) were stronger in the right hemisphere, the effect of the weaker alpha increase in the manipulation task appeared only in the left hemisphere (Hemisphere x TimeInt x Task interaction). In the left hemisphere the main effect of Task (i.e. Manipulation alpha < Retention alpha) was found during Delay1 (t(155) = 3.82, p < 0.001, dz = 0.31) and Delay2 (t(155) = 3.558, p < 0.001, dz = 0.33). In order to test whether the effect of Task was stable during the delay, we conducted another ANOVA using only the data in the left hemisphere and the delay period. No significant interaction TimeInt x Task was found (F(1, 155) = 1.27, p = 0.262, η^2^ < 0.01). To additionally test the hypothesis about the lack of time related changes in the effect of Task, we conducted a Bayesian ANOVA using the BayesFactor package for *R* with default priors. The resulting BF_01_ = 7.3 for TimeInt x Task interaction suggests that the model without the interaction is 7.3 times better supported by the data than the model with the interaction. Thus, the stronger suppression of alpha in the manipulation task as compared with the retention task is stable over time (see Figure 3c).

### Beta

The beta rhythm decreased during the task as compared with baseline, and this decrease was stronger in the manipulation task than in the retention one (main effect of Task, see Figure 4 and Table 4). Beta was stronger suppressed during Encoding than during Delay, and stronger during Delay2 than Delay1 (TimeInt main effect). Pairwise comparisons revealed strong effects distinguishing each level of TimeInt factor (Encoding vs Delay1: t(155) = 16.3, p < 0.001, dz = 1.31; Encoding vs Delay 2: t(155) = 11.8, p < 0.001, dz = 0.95; Delay1 vs Delay2: t(155) = 7.47, p < 0.001, dz = 0.6). Subsequent t-tests to examine the TimeInt x Task interaction showed that beta was stronger suppressed in the manipulation task only during Delay1 (t(155) = 4.9, p < 0.001, dz = 0.39) but not during Encoding (t(155) = 0.94, p = 0.38, dz = 0.08) or Delay2 (t(155) = 2.81, p = 0.011, dz = 0.23 after Holm’s correction).

**Table 4.** Beta ANOVA RM

| | *DF* | *F* | $\eta^2$ | *p* |
| --- | --- | --- | --- | --- |
| **TimeInt** | 1, 224 | 182.38 | 0.541 | **< 0.001 \*\*\*** |
| **Task** | 1, 155 | 11.34 | 0.068 | **0.001 \*\*** |
| Load | 2, 306 | 0.19 | 0.001 | 0.827 |
| **TimeInt x Task** | 2, 285 | 23.85 | 0.133 | **< 0.001 \*\*** |
| TimeInt x Load | 4, 580 | 0.63 | 0.004 | 0.628 |
| Task x Load | 2, 299 | 5.42 | 0.034 | 0.0055 |
| TimeInt x Task x Load | 4, 579 | 1.48 | 0.009 | 0.210 |

\* small \*\* medium \*\*\* large effect size and $p < 0.005$

The Task x Load interaction indicated possibly different load-dependent dynamics in the Manipulation and Retention tasks (see Figure 4). Taking into account the lacking effects of Load in all previous comparisons, it was worth investigating this formally non-significant (p = 0.0055) interaction. Two RM ANOVAs with the factor Load revealed no significant effects (manipulation task: F(2, 310) = 3.34, p = 0.037, η^2^ = 0.02; retention task: F(2, 296) = 1.98, p = 0.142, η^2^ = 0.01). Additionally, in order to check whether there is any evidence on the presence of the load effect at any time point, in any task and frequency band, we run cluster-based permutation tests separately on the manipulation and retention tasks data and three frequency bands. The tests yielded no significant clusters.

## Discussion

### Retention and Manipulation

#### Theta

Frontal midline theta activity was generally stronger during the task than in baseline. The role of theta activity in WM is hypothesized to be related to the maintenance of temporal relationship between items in memory (Hsieh & Ranganath, 2014). In previous studies, the requirement to keep the temporal order of events led to enhanced theta activity (Hsieh et al., 2011; Roberts et al., 2013). In the present study, the temporal order also played an important role in successful performance of both tasks. Even though the stimuli were presented simultaneously, they were unlikely to be rehearsed as a single visual pattern. Multiple studies convincingly showed that verbal information is spontaneously coded as ordered representations in WM along the horizontal axis (Gmeindl, Walsh, & Courtney, 2011; Guida, Abrahamse, & Dijck, 2020; Oleron & Tardieu, 1982). In the same vein, Okuhata et al. (2012) demonstrated a lack of difference in performance in a direct comparison of simultaneous and successive versions of the Sternberg paradigm. In our WM paradigm probing spatio-temporal order, we also observed a strong theta enhancement during delay (one-sample t(155) = 8.88, p < 0.001, Cohen’s dz = 0.71).

According to Hsieh & Ranganath (2014) the increase of the set-size would increase the complexity of temporal relationships between memory items. Therefore, not only the requirement of manipulation but also increasing memory load should result in theta enhancement during delay. However, no load-dependent effect was found. Theta increase with increasing WM load is not always a reproducible phenomenon (for review, see Pavlov & Kotchoubey, 2021). The reasons for the lack of the load effect are not clear. As a possible explanation, because all load levels used in the present study were rather high, theta could have already attained a plateau at the lowest level of load.

Theta activity was greater in the manipulation as compared to the retention task. The close relationship between manipulations in verbal WM and theta rhythm enhancement has been repeatedly observed in the literature. In similar studies with alphabetical reordering task the effect of enhanced theta in the manipulation condition has been replicated consistently (Berger et al., 2014; Griesmayr et al., 2010; Pavlov & Kotchoubey, 2017). The same effect was also found for other types of verbal WM manipulation (Itthipuripat et al., 2013; Kawasaki et al., 2010). Although in a recent spatial WM study by Berger et al. (2019) theta activity did not differ between the manipulation and retention tasks, it did in three other studies using the same experimental paradigm (Berger et al., 2016; Eschmann, Bader, & Mecklinger, 2018; Griesmayr et al., 2014). Thus, as an addition to the previous research, we showed that stronger theta enhancement in more executive control demanding WM tasks is a replicable phenomenon at least in the verbal domain.

One might argue that the manipulation task was simply more effortful. From this point of view, not the increased involvement of the executive control networks but a general task difficulty (i.e., the required effort) produced the theta effect. However, the most obvious measure of task difficulty is the performance. The performance, as measured by the accuracy, was similar in Manipulation of 5 and Retention of 7 letters conditions (mean±SD: 79.55±14.33 and 80.71±13.55 in Manipulation 5 and Retention 7 conditions, respectively). Nevertheless, the difference in theta spectral power between these two conditions was still substantial (main effect of Condition: F(1, 155) = 10.55, p = 0.0014, η^2^ = 0.06).

The difference between the tasks in the power of the theta frequency band was not stable in time. Theta started differentiating between the tasks at about 1 s in the encoding period and the difference disappeared at around 4 s in the delay period. This temporal profile may reflect the trajectory of the alphabetizing process. In support of this claim, the time to retrieve items from a memorized sequence of 5 letters was at least 4 seconds for the first item and less than a second for the following items (Salthouse & Coon, 1993). For 6 and 7 letter sets the number can be larger. We can assume that (some) participants probably started manipulations while the stimuli were still on the screen and continued the task until it is completed – as expected, in the middle of the delay period. This finding is another important piece of evidence confirming that theta power increase reflects the increased demand on executive control network to comply with the requirement of the task.

#### Alpha

Generally, posterior alpha activity showed a pattern of suppression during encoding that was reversed during the delay in both tasks. The pattern of alpha suppression during encoding, which switched to a continuous enhancement starting from 1 s after the delay onset, has been shown before (Embury et al., 2018, 2019; Heinrichs-Graham & Wilson, 2015; McDermott et al., 2016; Proskovec, Heinrichs-Graham, & Wilson, 2016; Wiesman et al., 2016; Wilson et al., 2017), although contradicting examples with the opposite pattern (i.e., alpha suppression during delay) are not rare (Pavlov & Kotchoubey, 2021).

Previous research has attributed the alpha suppression to the cortical engagement allowing either encoding of information into WM or decoding the information for retrieval (Jensen & Mazaheri, 2010; Klimesch, Sauseng, & Hanslmayr, 2007). On the other hand, the alpha increase during delay may reflect sensory gating through the disengagement of certain cortical areas to protect memory representations from interference (Klimesch et al., 2007; Payne & Sekuler, 2014; Roux & Uhlhaas, 2014). Thus, alpha suppression during encoding in our study probably reflects the active state of visual information processing. The alpha enhancement at the start of delay signalizes switching attention to the internal representations supported by blocking of visual input to counteract possible interference.

On average, alpha returned to the baseline level around the middle of the delay period. Nevertheless, the results showed that the effect of stronger alpha suppression in the manipulation task was present throughout the whole delay period. The difference between the tasks was not observed during encoding. The decrease of alpha during delay in the manipulation task suggests weaker inhibition of cortical areas actively involved in the processing of the information. A possible explanation is that the manipulation task requires continuous recoding of the information; therefore, the encoding processes are not finished by the beginning of the delay period but continue. However, the difference between manipulation and retention was observed even at the end of the delay when mental manipulations are expected to come to an end.

The effect of stronger alpha suppression in the manipulation than in the retention task was present only in the left hemisphere, possibly indicating involvement of the language cortex. One might object to this hypothesis, that this involvement should also be manifested in an effect of WM load, but this effect was not found. However, it is plausible that the difference (in terms of engagement of the language cortex) between the levels of load at the rather high range from 5 to 7 items is much subtler than the difference between the types of the task.

The above mentioned effects are suggestive of the idea that mapping the patterns of alpha suppression/enhancement on a single process is counterproductive. The alpha dynamics during performance of a WM task is rather a combination of multiple tendencies such as general level of activation, visual perception block by cortical disengagement and most likely other processes.

#### Beta

Unlike theta and alpha, centrally localized beta activity was inhibited throughout the task as compared with baseline. In the first part of the delay period, beta was more strongly suppressed in the manipulation than in the retention task. In contrast to alpha and theta, the literature does not shed any light on the possible functional role of beta oscillations in WM. Our results are in line with the only available verbal WM study comparing the components of WM (Berger et al., 2014). Berger et al. argued for the status quo model of beta activity that asserts that beta oscillations are related to the maintenance of the current cognitive state (Engel and Fries, 2010). According to the model, beta is high when no changes in the cognitive state appears, and suppressed as soon as new information processing is required. Updating and manipulating information in WM would change the current state of memory thus suppressing beta activity. The manipulation task required constant attention switching between the modified reordered string of letters and the original one. Above we already hypothesized that the reorganization of memory should be finished by the second half of the delay period, which can be regarded as the establishment of a stable “status quo”. This is exactly the time when the difference in beta activity between manipulation and retention disappeared. This finding is in line with the status quo model.

Against the model, however, speaks the fact that the average power of beta in the second part of the delay was lower than in the first part of it, while the model predicts the opposite relation. However, the beta suppression in the second part of the delay can be explained by preparation of the motor response. Similar beta activity between 15-20 Hz was observed by Proskovec, Heinrichs-Graham, et al. (2019) in the Sternberg task. Their beta appeared about 800 ms before the probe. Preparation to the following motor response might lead to such activation. However, response preparation alone is not a satisfactory explanation for the beta effects obtained in the current experiment, because the effect of task in beta frequency band was significant only in the first part of the delay when no motor preparation is meaningful. Furthermore, motor preparation during encoding cannot explain even larger beta suppression than during delay period.

Another thinkable explanation might be that the motor cortex is involved not only in motor response preparation but also in unintentional (imaginary) movements. The mental manipulations in the present task involve a replacement of imaginary objects (letters). This operation may engage motor areas related to the control of hand movement and areas such as frontal eye fields (FEF) related to the control of eye movements. FEF are a part of a dorsal attention network heavily involved in the process of encoding of information to WM (Kim, 2019). Contrary to this hypothesis, a review of fMRI studies failed to find any involvement of premotor areas in WM manipulation tasks (D’Esposito, Postle, & Rypma, 2000).

A possible kind of motor activity occurring in WM tasks is subvocal rehearsal of the presented items. In fact, many participants reported that they used this strategy, but its influence on EEG oscillations is difficult to evaluate due to extreme scarcity of experimental evidence. Plaska, Ng, & Ellmore (2021) compared maintenance of two images with an articulatory suppression and with standard rehearsal of verbal labels assigned to the images. The authors associated rehearsal processes with the enhancement of alpha and beta activity in the first half of the delay and its subsequent suppression in later stages (after about 4 s). Another study compared verbal and nonverbal content memory and found an increase in oscillatory brain activity probably related to subvocalization practically in all frequency bands (Hwang et al., 2005). The beta band demonstrated the most consistent effects, particularly over fronto-central and parietal areas, pending the authors to relate beta activity to subvocal rehearsal. Although alpha and beta activity can be related to subvocal rehearsal, this fact fails to provide any explanation of the observed patterns in our data.

To sum up, three possible factors may determine the dynamics of beta oscillations: the involvement of subthreshold motor activity such as subvocalization or activation of premotor cortical areas during mental manipulations, the maintenance of status quo (Engel & Fries, 2010) after finishing the manipulations, and response preparation at the end of the delay interval.

### Duration of the delay

A long delay period was employed to investigate the effect of its duration on the oscillatory brain activity. In the literature, there is no evidence of verbal WM performance decline with increased duration of the delay when no concurrent task is present (Oberauer, Farrell, Jarrold, & Lewandowsky, 2016; Oberauer et al., 2018). Therefore, one may expect to observe constant neural activity throughout the delay period. Contrary to this expectation, oscillatory brain activity strongly varied across the delay period in all studied frequency bands, thus indicating the complexity of processes taking place during this time.

Prominently, alpha activity faded out in the second part of the delay. Heinrichs-Graham & Wilson (2015) suggested that at later stages of WM tasks the alpha activity is required to counteract forgetting. This hypothesis, however, is not supported by the present data demonstrating that at the end of a long delay period, alpha returns to the baseline level despite the increasing likelihood of forgetting with time. In all previously mentioned studies, this phenomenon could not be found because the delay was only 3 s long. Ellmore et al. (2017) is the only available WM study with a long enough delay (6 s) that reported TF representations of alpha activity and a formal statistical analysis of the time effect. The authors also demonstrated that alpha activity behavior during the delay was transient rather than sustained.

There is mounting evidence that information in WM can be maintained without detectable neural activity related to the maintained information. Strong support for this idea comes from machine learning studies aiming to decode the content of WM from EEG and fMRI data (Bae & Luck, 2018; Lewis-Peacock, Drysdale, Oberauer, & Postle, 2011; Rose et al., 2016; Wolff, Jochim, Akyürek, & Stokes, 2017). The studies showed the possibility to match neural patterns with specific content currently maintained in WM. Thus, Wolff et al. (2017) used lateralized alpha-activity in the retro-cue paradigm as a read-out of WM content. In this paradigm, participants had to encode two items located on both sides of the fixation point. A retro-cue, presented after a short delay, informed the participants which item should be kept in memory and which item is no longer relevant. The alpha activity demonstrated a lateralized pattern being stronger suppressed over the hemisphere contralateral to the cued item. The authors were able to decode above the chance level the content of WM until the end of the delay. However, the figures show that decoding accuracy quickly deteriorated and even approached the chance level towards the end of delay (<1 s). In another study employing the same approach the accuracy of the classification declined below chance level shortly before the end of delay, i.e., 700 ms after the delay onset (Wolff, Kandemir, Stokes, & Akyürek, 2019). The fact that it was possible to decode only the attended item, led to a conclusion that neural activity may represent not the entire content but only information in the focus of attention (Lewis-Peacock et al., 2011; Stokes, 2015; Wolff, Ding, Myers, & Stokes, 2015; Wolff et al., 2017). Interestingly, the unattended information lost decodability even faster. Nevertheless, all the information, whether decoded or not, was remembered after the end of delay.

The information that is still available for retrieval but not detectable through correlates of neural activity was termed as activity-silent (Stokes, 2015). The inability to decode information from such a noisy signal as the EEG does not provide strong evidence for the activity-silent model of WM because it may be related to the limitations of the method. However, the data of intracranial recordings support the notion that the inability to detect WM content in EEG cannot be attributed only to the technical limitations. In monkeys, the neural activity related to WM maintenance can completely disappear during the delay returning only shortly before the probe when a decision has to be made (Barak, Tsodyks, & Romo, 2010). Typical WM experiments in monkeys use only one item for encoding, but in a study employing a Sternberg-type multi-item visual WM task (Konecky, Smith, & Olson, 2017), only the last item of a multi-item sequence was represented by persistent firing while the other items were activity-silent. The activity-silent model of WM suggests that rapid changes in synaptic weights allow to maintain information in WM even in the absence of persistent neural activity (Manohar, Zokaei, Fallon, Vogels, & Husain, 2019; Silvanto, 2017; Stokes, 2015). Probably, persistent activity maintains the memory when the remembered item is in the active state of focused attention. When attention is shifted to another item, unattended WM items are stored in activity-silent synaptic traces.

An alternative model postulates a separation of the processes taking place during the WM delay period into iconic, transient and sustained stores (Ruchkin, Grafman, Cameron, & Berndt, 2003). The stores are characterized by timing of the activation. The transient store in the model operates over about 4 s after the delay onset. The role of the transient store is to translate information from iconic into a more sustained long-term form. Following this view, the actual duration of what we call WM or short-term memory is about 4-5 seconds long, after which long-term memory (LTM) begins. Corroborating the idea, performance in WM shows a similar deficit after about 5 s like in LTM tasks in patients with medial temporal lobe lesions (Jeneson & Squire, 2011). The first 4 s may represent a sensitive period of memory consolidation. Ranganath et al. (2005) showed that presentation of a visual distractor after 1 s of delay disturbed LTM performance, whereas a presentation of the distractor 4 s after the delay onset did not affect later recall. These findings suggest intriguing similarities between WM after 4 s delay and LTM.

We observed neither persistent neural delay activity nor completely activity-silent delay. Rather, the activity strongly varied along the delay period. Although our data cannot be conclusive regarding the relationship between WM and cortical oscillations, we putatively hypothesize that the resulting pattern may be determined by an interplay of two mechanisms. At the beginning, neural firing is the leading mechanism supporting memory trace. As the time passes, the weight of this mechanism decreases, and another mechanism of rapid plasticity becomes more important. Perhaps, the latter mechanism fully takes over the responsibility of maintaining information in WM after about 4 seconds of the delay. At present this hypothesis remains speculative, and more research is needed to test it.

## Conclusions

In the current exploratory analyses of our publicly available dataset, we confirmed the results of previous studies showing that the oscillatory EEG activity strongly differs between the manipulation and retention tasks. This difference had diverse temporal trajectories. It maintained over encoding and the first part of delay in theta, during the first part of delay in beta, and during the whole delay period in alpha. Assuming that the increased theta power reflects involvement of executive functions, the present data suggest that the executive control of attention started already during the encoding period. Alpha, in turn, was likely to reflect the activation of language areas and sensory input blockade during delay; finally, the power of beta was related to the mental manipulations when visual input is blocked.

## Author contributions statement

YGP: Conceptualization, Funding acquisition, Investigation, Data curation, Formal analysis, Project administration, Visualization, Methodology, Writing - original draft, Writing - review & editing.

BK: Writing - review & editing.

## Data availability statement

The data are publicly available on OpenNeuro (doi:10.18112/openneuro.ds003655.v1.0.0).

## Acknowledgements

Study was supported by Russian Foundation for Basic Research (RFBR) #19-013-00027.

